# Disrupting FOXO4 function confers neuroprotection against oxidative stress and ischemia-reperfusion-caused neuronal injury

**DOI:** 10.1101/2025.03.13.643180

**Authors:** Yasin Asadi, Farhad Gorjipour, Rozenn K. Moundounga, Abena Dwamena, Augustina Potokiri, Erin Gilstrap, Xiaoping Li, Caroline McDowell, Hongmin Wang

## Abstract

Previous data suggest that FOXO4 facilitates inflammation and oxidative stress in *non-brain tissues* under stress or disease conditions, indicating that blocking FOXO4 function may be neuroprotective in ischemia-reperfusion-induced brain injury. However, this possibility has not been tested in a cerebral ischemia-reperfusion condition. Here, we treated the FOXO4 knockout (KO) primary neuronal cultures with oxidative stress or oxygen-glucose deprivation (OGD) or subjected the KO mice to the transient middle cerebral artery occlusion (tMCAO). Our results showed that KO of FOXO4 reduced oxidative stress and OGD-induced neuronal death, attenuated tMCAO-caused infarct volume, improved animal survival, decreased neurological deficits, and enhanced functional recovery compared to the WT cells or mice. Immunohistochemical staining and Western blot analysis suggested decreased neuroinflammation in the KO brain. These data indicate that FOXO4 is a therapeutic target, and disrupting its activity promotes neuronal survival following ischemic stroke-induced brain injury.

## Introduction

Stroke is a major health problem: it is the fifth leading cause of death and the leading cause of disability in the United States. Ischemic stroke is the most frequent type of stroke, affecting millions of people across the globe (1). Despite intensive research, there is still no effective treatment for this condition, and tissue plasminogen activator remains the only Food and Drug Administration-approved medication for treating ischemic stroke (2). While it is necessary to restore cerebral blood flow, reperfusion itself induces the production of large amounts of reactive oxygen species that further damage proteins and other intracellular macromolecules to exacerbate brain injury (3). Moreover, ischemia/reperfusion (I/R) also disrupts the blood-brain barrier, resulting in infiltration of leukocytes and inflammatory responses to aggravate brain injury (4). To develop effective therapeutics, it is necessary to identify and validate additional therapeutic targets.

Transcriptional gene regulation, governed primarily by numerous transcriptional factors, is one of the upstream parts potentially modulating I/R-induced brain injury. The Forkhead Box Proteins O (FOXOs), consisting of four structurally and functionally related proteins, FOXO1 (also referred to as FKHR), FOXO3 (also known as FOXO3a), FOXO4 (also known as AFX1), and FOXO6, are one family of transcription factors, representing mammalian homologs of daf-16 in *C. elegans* and playing a crucial role in cell survival, cell proliferation, metabolism, response to oxidative stress, apoptosis, and aging (5). Under oxidative stress or the absence of the cellular survival drive of growth factors, FOXOs translocate to the nucleus and upregulate a series of target genes, thereby promoting cell growth arrest and apoptosis (6, 7).

Despite the highly structural and functional similarities of FOXOs, previous data have suggested that the physiological roles of FOXOs are functionally diverse in mammals (8, 9). For instance, FOXO1 deficiency is embryonically lethal, and FOXO3 deficiency exhibits age-dependent infertility in females in mice; however, the loss of FOXO4 does not cause any notable changes in the mouse, suggesting that FOXO4 may play a different role from other FOXO members (8, 10). FOXO4 was found to promote cell death in the heart and the liver following I/R, while downregulation of FOXO4 confers protection against I/R-induced tissue injuries in these organs (11, 12). Moreover, following cardiac ischemia, FOXO4 enhances the interaction of leukocytes with the endothelial cells of blood vessels to promote early tissue inflammation (13). In contrast, downregulation of FOXO4 suppresses oxidative stress-induced cell death in proangiogenic cells and promotes neovascularization in ischemic limbs (14). Functionally, FOXO activity is negatively regulated by phosphorylation via the phosphoinositide 3-kinase-Akt pathway, a well-known cell survival pathway (6, 7). Ischemic preconditioning upregulates Akt activity, leading to FOXO inhibition and promotion of neuronal survival against a subsequent severe ischemic insult (15). These prior studies suggest that FOXO4 may be a therapeutic target for ischemic stroke, yet this has not been tested in ischemic stroke neuronal cells or animals. In this study, we examined the role of FOXO4 in neuronal injury *in vitro* and *in vivo* following oxidative stress, oxygen-glucose deprivation (OGD), and I/R, using the primary neuronal cultures and FOXO4 knockout (KO) mice combined with different approaches.

## Materials and Methods

### Mice

All animal-related experiments and procedures were approved by the Institutional Animal Care and Use Committee of the Texas Tech University Health Sciences Center and were in compliance with the National Institute of Health Guide for the Care and Use of Laboratory Animals. Animals were maintained on a 14-/10-hour light/dark cycle with a maximum of four mice per cage, able to access water and food ad libitum. FOXO4 global knockout (KO) mice in the background of FVB have been described before (13) and were used for ischemic stroke-induced brain infarct studies, while the same strain of wild-type (WT) mice was used as the control. For animal behavioral studies, the KO and WT male mice with a hybrid background (F1 generation) produced from the heterozygous KO FVB females crossed with the WT C57BL/6 males were used, because of the retina degeneration problem in the FVB mice. As FOXO4 is an X chromosome-linked gene, in this breeding strategy, 50% of males would be FOXO4 KO males, and 50% would be WT males that were used as controls for behavioral studies following ischemic stroke. More than one litter of animals was used in each experiment. The male animals between 8–12 weeks of age with a body weight of 25-30 g were used in the study. Sample size calculations and power analysis were performed according to our previously described methods (16) using the statistical software Stata (StataCorp LP, College Station, TX, USA).

### Primary cortical neuronal cultures

Primary cortical neuronal cultures were prepared from wild-type and FOXO4 KO mice at postnatal day 0 according to our previously described methods.(17) Briefly, the cerebral cortex was isolated and digested with 5 ml of 0.25% trypsin/EDTA supplemented with 75 µl 0.1% DNase (2000 UI/mg). After 15-20 minutes of digestion at 37°C, the digested tissues were centrifuged at 1000 × g for 3 minutes at room temperature. The supernatant was discarded, and the resulting tissue pellets were resuspended with fetal bovine serum (FBS) to stop the digestion. After centrifugation, the tissues were resuspended with primary neuronal culture medium (Neurobasal medium supplemented with 2% B27, 2 mM L-glutamine, and penicillin/streptomycin) and then pipetted up and down several times using serological pipets to disrupt tissues. After filtering through a 70 μm nylon cell strainer, the cell suspension was adjusted and plated in poly-DL-lysine-coated 12-well plates. After 7 days of incubation at 37°C in a 5% CO2 incubator, the cells were treated with 20 µM menadione (MD) for 24 h or oxygen-glucose deprivation (OGD) for 3 h and then in normal culture conditions for 21 h before cell viability was assessed.

### Oxygen-glucose deprivation (OGD)

WT and FOXO4 KO primary neuronal cultures were subjected to OGD at 7-10 *in vitro* days according to previous methods (17). Briefly, the neuronal culture medium was replaced with a glucose-free Hank’s Balanced Salt Solution (HBSS), and then the culture plates were placed into a hypoxic chamber where the air was replaced with a 5% N_2_ and 95% CO_2_. After 3 h of OGD condition, the neuronal cultures were returned to the normal culture condition, and after 21 h cell viability was measured via ATP and MTT assays.

### ATP assay

We used an ATP detection kit (Cayman Chemical, Ann Arbor, Michigan, USA) to measure ATP levels in cultured neurons according to the manufacturer’s instructions.

### MTT assay

Cell viability in cell cultures was assessed using an MTT assay kit (R&D Systems, Minneapolis, MN, USA) based on the company’s guide.

### Transient middle cerebral artery occlusion (tMCAO)

The tMCAO procedure was performed according to previously described methods (18, 19). Briefly, anesthesia of mice was induced with 5% isoflurane and then maintained with 2% isoflurane. The left hemisphere was subjected to tMCAO using a silicon-coated monofilament (RWD, Sugar Land, TX, USA), and after 1 hour, the monofilament was removed. A Laser Speckle Imaging System (RWD, Sugar Land, TX, USA) was used to monitor cerebral blood flow through the MCA. Only the mice with successful occlusion of MCA were included in the studies, as reflected by reduced blood flow by over 80% and reperfusion with more than 75% recovery of blood flow in the MCA.

### TTC staining and measurement of infarct volumes

Mice were euthanized 24 h after I/R to isolate the brains. After being sectioned into 2 mm slices, the brain sections were incubated with 2% of 3,5-triphenyl tetrazolium chloride (TTC, Sigma, Saint Louis, MO, USA) at 37°C for 20 minutes. Subsequently, the sections were fixed in 4% paraformaldehyde and then imaged. The infarcted volume was measured using Image J software and calculated as previously described (20).

### Assessment of modified neurological deficits

At 1, 3, 5, and 7 days following I/R, a modified neurological severity score (mNSS) system was applied to assess motor and sensory functions in mice (21). Based on this system, a scale of 0 to 14 was given to an animal, with 0 for normal neurological behavior and 14 for the maximal neurological deficit (22).

### Novel object recognition (NOR) test

A box (50 × 50 × 30 cm) containing two objects was used for the NOR test at 7 days following I/R. On the first day of the test, mice were placed in the box and allowed to move freely for 10 minutes. On the second day, one of the objects was replaced with a new one (novel object), and the other remained as a familiar object. Then, mice were placed into the box to move and explore the objects. By using a camera, the total time spent exploring each object was recorded, and the ratio of the new object exploration was calculated as the discrimination index (DI = (Tnovel - Tfamiliar)/(Tnovel + Tfamiliar) (19).

### Immunofluorescent staining

Brains were coronally cut into 3 parts, and the frontal and occipital parts were used for TTC staining to define the penumbral area. The middle part was washed and fixed in 4% paraformaldehyde. Then, the 16 μm thickness sections were prepared on a Cryostat (Leica, Buffalo Grove, IL, USA), and the brain sections were processed based on the prior method (23). The primary antibodies used were anti-GFAP (1:50, EMD Millipore, #MAB360), anti-Iba1 antibody (1:50, Cell Signaling Technology, #31659), anti-CD45 (Invitrogen #MA180090), anti-Ly6G (Cell Signaling #87048), and anti-CD11 (Developmental Studies Hybridoma # M1/70.15.11.5.2). The secondary antibodies were Texas red-conjugated anti-rabbit and FITC-conjugated anti-mouse antibodies. DAPI (IHC-tek#IW-1404) was used to stain the nuclei.

The images were captured with a fluorescence microscope (Echo Revolve, San Diego, USA) and quantified with ImageJ software (24). The ratio of the red- or green-positively stained cells in each field was calculated by the total red or green cells divided by the total number of cells (DAPI positively stained cells) (25, 26).

### Western blot

The cortex of the left (i.e., the I/R side) hemisphere of each mouse brain was collected for lysate preparation in the RIPA buffer supplemented with a protease cocktail. The brain tissues were sonicated in the lysate buffer, and the lysates were centrifuged at 12,000 x g for 4 min at 4°C. The supernatant was collected, and then the total protein concentration was measured before being subjected to the SDS-PAGE and transferred to the nitrocellulose membrane according to previous methods (17). The antibodies used were anti-IL-1β (1:1000, Cell Signaling, #63124), anti-IL-6 (1:1000, Cell Signaling, #12912), anti-TNFα (1:1000, Cell Signaling, #11948), and anti-β-actin (1:1000, Santa Cruz Biotechnology, #sc-1616). The secondary antibodies used were anti-rabbit IgG, HRP-conjugated antibody (Cell Signaling, #7074), and anti-mouse IgG, HRP-conjugated antibody (Cell Signaling, #7076). The western blot results were documented using an imaging system (C400 Azure Biosystem), and protein band intensities were measured using the NIH ImageJ software.

### Statistical analysis

Statistical analyses were conducted using GraphPad Prism version 9.0 statistical software. Differences between the two groups were assessed using an unpaired t-test. For comparisons between more than two groups of animals, repeated measures and two-way ANOVA with Tukey’s post hoc test were utilized. All numerical data were presented as mean ± SD or SEM. P < 0.05 was regarded as statistically significant.

## Results

### KO of FOXO4 reduces neuronal death caused by oxidative stress or OGD in primary cortical neuronal cultures

To assess the effect of KO of FOXO4 on oxidative stress-induced cell death, we treated the primary cortical neuronal cultures with an oxidative stress inducer, menadione (MD)(18), and then measured the viability of the cells via ATP and MTT assays. KO of FOXO4 was confirmed by Western blot analysis of FOXO protein level (**Fig. 1a**). Following MD treatment, KO neurons showed increased cell survival as reflected by higher ATP levels in KO cells compared to WT cells (**Fig. 1b**). Increased viability in the KO cells was also supported by the MTT assay (**Fig. 1c**). However, in those neuronal cultures without MD treatment, the cellular viability did not differ between WT and KO neurons (**Fig.1b, 1c**). To validate these results, we further treated the two types of neuronal cultures with an *in vitro* model of ischemic stroke, OGD, and then examined cell viability. In both the ATP assay (**Fig. 1d**) and MTT assay (**Fig. 1e**), KO neurons showed reduced cell death compared to WT neurons. These results indicate that KO of FOXO4 is neuroprotective against oxidative stress- or OGD-induced neuronal death in primary neuronal cultures.

**Figure 1.**
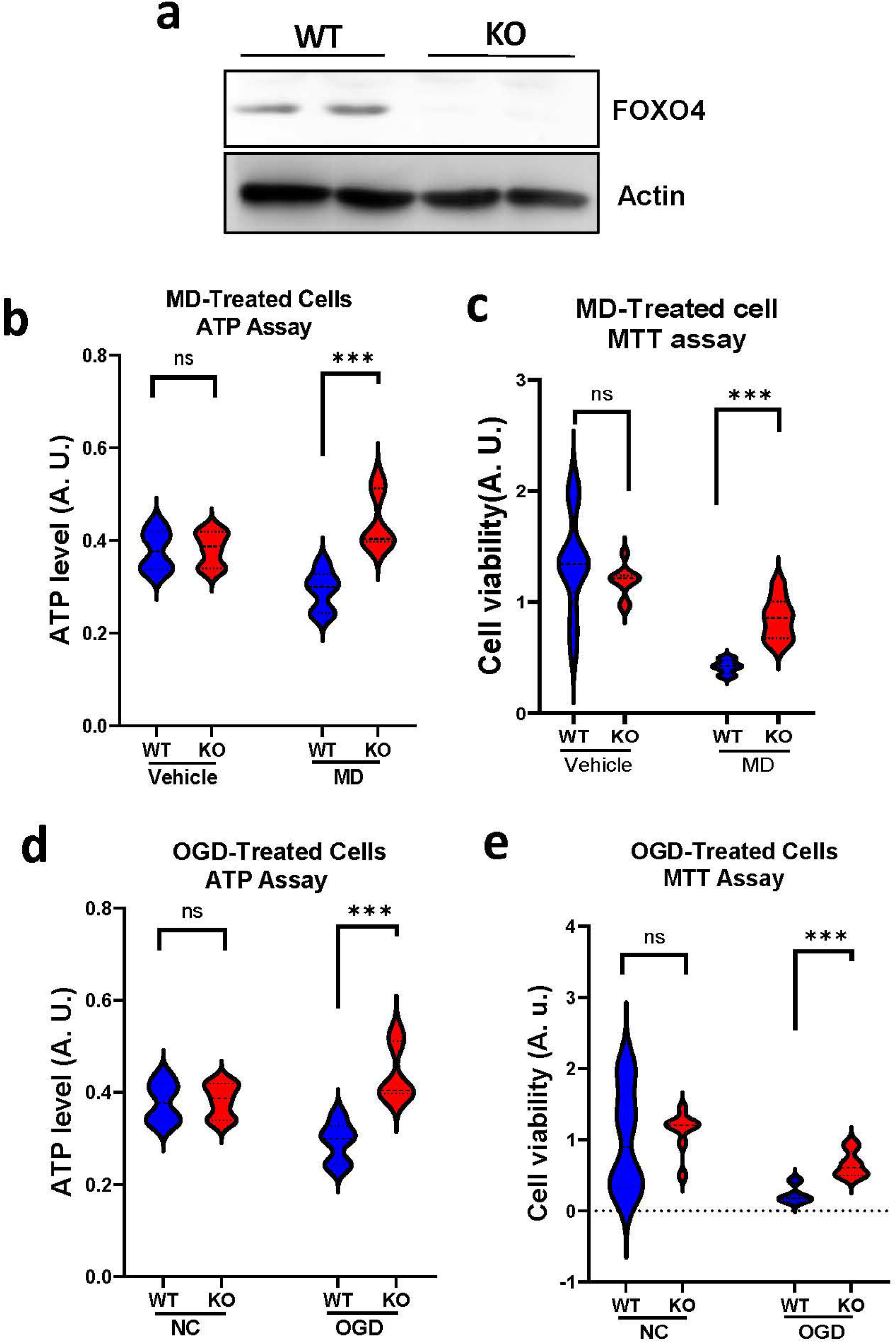
Effect of KO of FOXO4 on oxidative stress- and OGD-induced neuronal death in primary neuronal cultures. **a.** Western blot analysis of FOXO4 protein levels in primary neuronal cultures to validate FOXO4 KO. **b.** ATP assay results of KO and WT neurons following MD treatment. **c.** MTT assay results of KO and WT neurons following MD treatment. **d.** ATP assay results of KO and WT neurons following OGD treatment. **e.** MTT assay results of KO and WT neurons following OGD treatment. Data are shown as mean ± SD. N = 10-14, ***p < 0.001. ns, no significant difference.

### KO of FOXO4 reduces the infarct volume and enhances functional recovery after tMCAO

To determine whether FOXO4 influences the outcome of I/R-induced brain injury *in vivo*, we performed tMCAO in FOXO4 KO and WT mice. Selective KO of the gene was validated by Western blot analysis (**Fig. 2a**). After 24 h following the reperfusion, mice were euthanized to assess brain injury by the TTC staining (**Fig. 2b**). Our results showed a significant decrease in infarct volume in the KO group compared to the WT group (**Fig. 2c, 2d**), suggesting that disrupting the FOXO4 gene is neuroprotective against I/R-induced neuronal injury.

**Figure 2.**
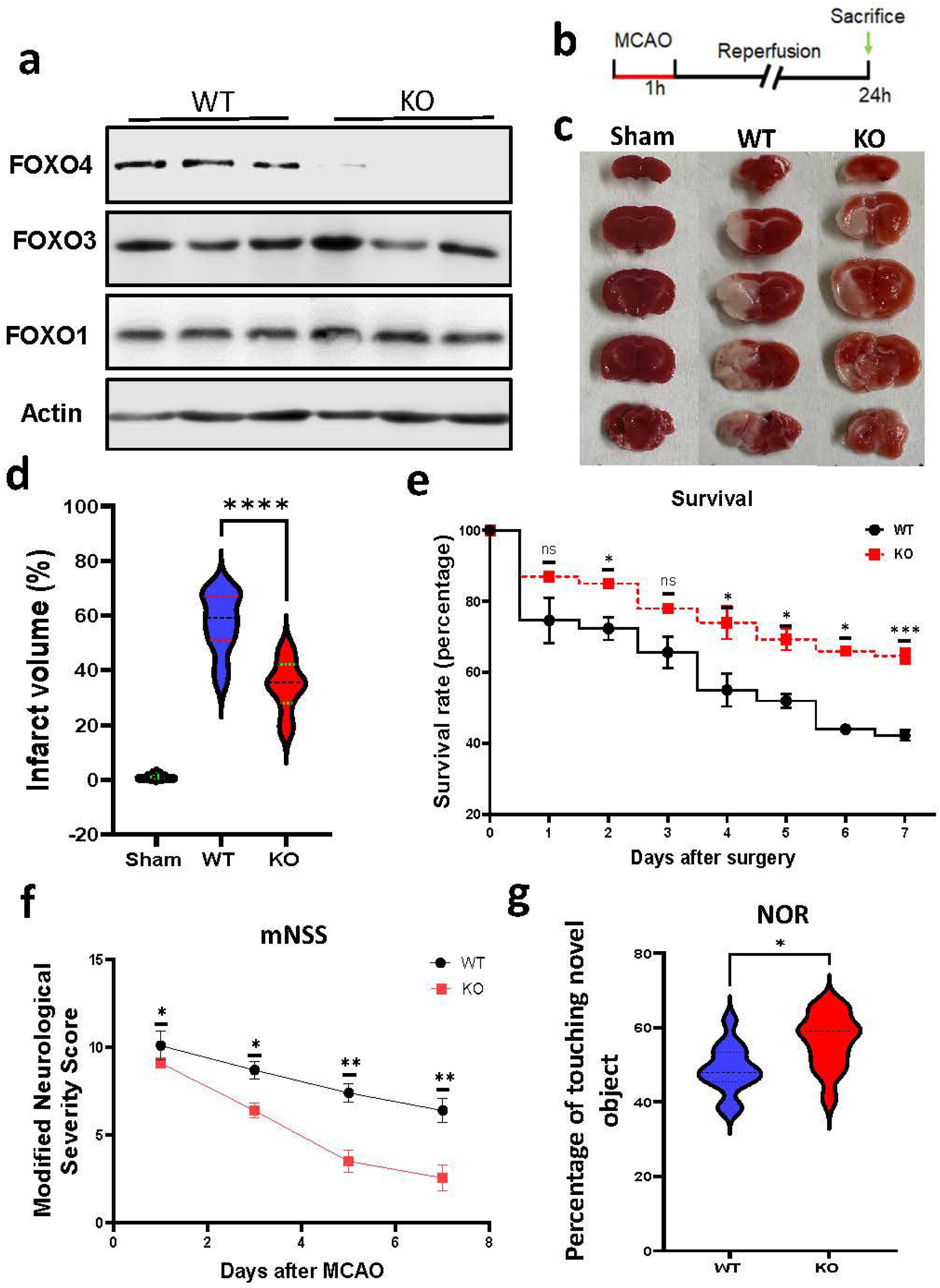
KO of FOXO4 reduces brain infarct volume, improves animal survival, and enhances functional recovery following I/R. **a.** Western blot analysis of FOXO4 protein levels in mouse brain to validate FOXO4 KO. **b.** A schematic of experimental design. **c.** Representative TTC staining of mouse brains from the sham surgery, WT, and KO animal groups after MCAO. **d.** Quantitation of TTC staining results. **e.** KO mice increased the animal survival rate compared to WT mice after tMCAO. **f.** KO mice improved mNSSs compared to WT mice after tMCAO. **g.** KO mice showed better memory compared to WT mice in the NOR test after tMCAO. Data are shown as mean ± SD. N = 10-14 for **(d)**, ****p < 0.0001. N = 14– 15 for (**e**)-(**g**), * p < 0.05, **p < 0.01, ***p < 0.001.

To determine whether KO of the FOXO4 influences animal survival following I/R, animal death and survival were recorded on days 1, 2, 3, 4, 5, and 7 after tMCAO. As shown in **Fig. 2e**, KO of FOXO4 increases the animal survival rate. Moreover, we also assessed the neurological deficits and found that KO of FOXO4 facilitated animal functional recovery, as reflected by the reduced mNSSs compared to the WT mice following the tMCAO procedure on days 1, 3,5, and 7 (**Fig. 2f**). Improved functional recovery in the KO mice was also supported by the cognitive function test, including the memory capabilities that were evaluated 10 days after MCAO. We observed better learning and memory in the KO mice compared to WT mice in the NOR test (**Fig. 2g**). Therefore, KO of FOXO4 reduces neuronal death, improves animal survival, and enhances functional recovery following I/R.

### KO of FOXO4 attenuates neuroinflammation in the brain following I/R

Neuroinflammation is a hallmark following I/R in the brain. To assess whether KO of FOXO4 alters the inflammatory responses, brain tissues on the ipsilateral side were isolated from KO and WT mice 48 h after I/R. Western blot analysis of the pro-inflammatory cytokines, including TNF-α, IL-1β, and IL-6, revealed a significant reduction in the KO brains compared to WT brains (**Fig. 3a-3d**). These data indicate that KO of FOXO4 attenuates I/R-induced neuroinflammation.

**Figure 3.**
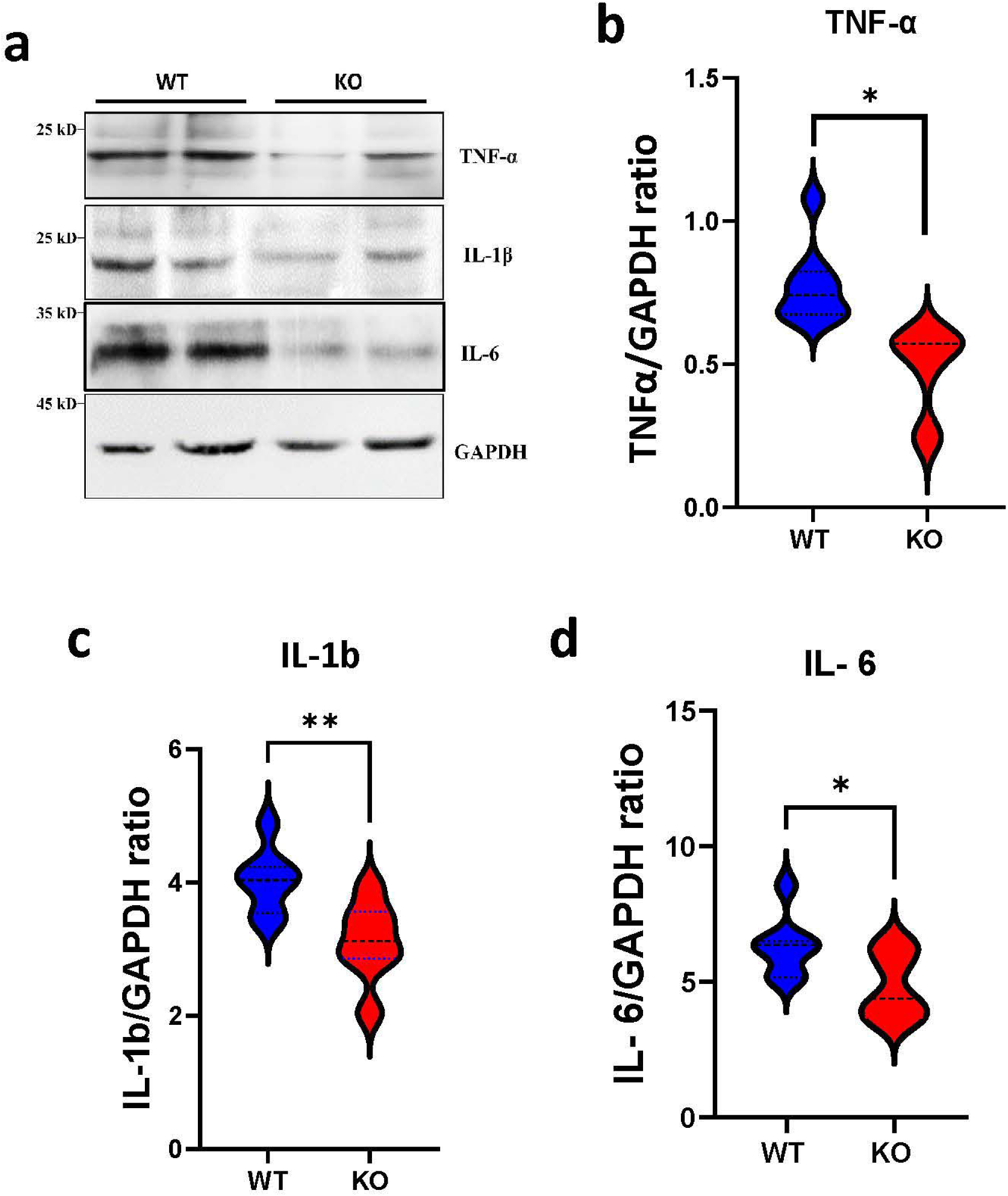
KO of FOXO4 attenuates neuroinflammation following I/R. **a.** Representative western blot results of the indicated proteins. **b-d.** Quantified results of the indicated proteins. Data are shown as mean ± SEM, * p < 0.05, **p < 0.01.

### KO of FOXO4 suppresses activation of astrocytes and microglia in the brain following I/R

To assess the reactive astrocytes and microglia, we performed immunostaining of the astrocyte marker, GFAP (glial fibrillary acidic protein), and microglia marker, Iba1 (ionized calcium-binding adaptor molecule 1), in the mouse brain 48 h following I/R. As shown in **Fig. 4**, the number of GFAP (**Fig. 4a, 4c**) and Iba1 (**Fig. 4b, 4d)** positively stained cells was significantly reduced in the KO brain compared to the WT brain, suggesting that the KO of FOXO4 suppresses gliosis following I/R in the brain.

**Figure 4.**
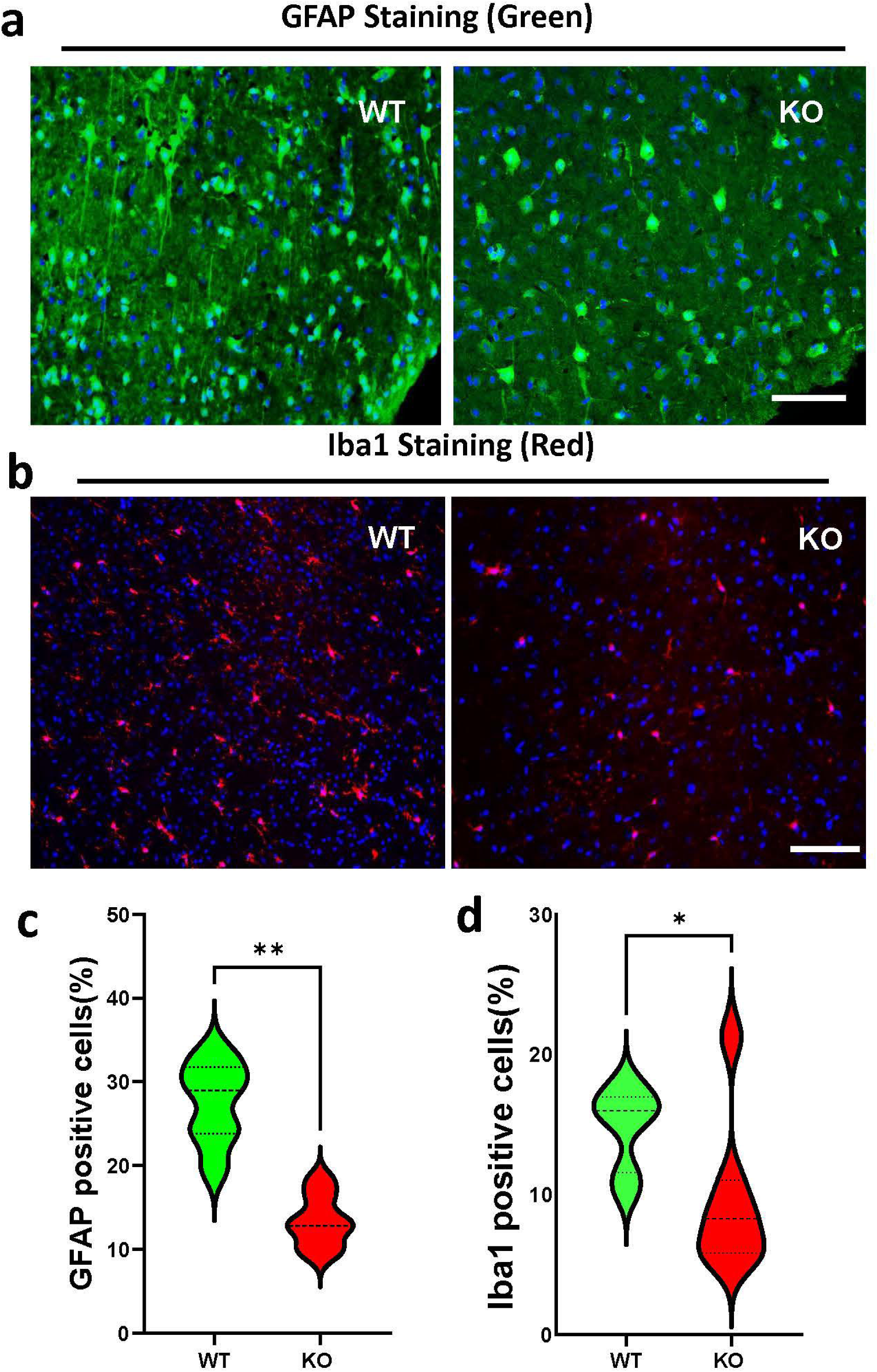
KO of FOXO4 reduces gliosis two days after I/R compared to WT mouse brains. **a.** Representative images showing GFAP staining in the WT and FOXO4 KO brains. Scale bar, 50 µm. **b.** Representative images showing Iba1 staining in the WT and FOXO4 KO brain sections following I/R. Scale bar, 50 µm. **c.** Quantitation of GFAP positively stained cells (green). **d.** Quantitation of Iba1 positively stained cells (red). All numeric data are shown as mean ± SD; n = 7 for each group. *p < 0.05, **p < 0.01.

### KO of FOXO4 reduces leukocyte infiltration in the brain following I/R

Another pathological feature following I/R is the infiltration of leukocytes into the brain. To determine the leukocyte infiltration, the brain sections were immunohistochemically stained either with the CD45 (a marker for the total hematopoietic cells), CD11b (a marker for macrophages, granulocytes, and NK cells), or Ly6G (a marker for monocytes, granulocytes, and neutrophils). As shown in **Fig. 5a** and **5b**, the number of the CD45 positively stained cells in the KO brain showed a reduction trend compared to the WT brain, despite the lack of statistical difference between the two groups. In contrast, the number of positively stained CD11b was significantly decreased (**Fig. 5c** and **5d**) in the KO brain compared to the WT brain. Similarly, the number of positively stained Ly6G (leukocytes) was also significantly decreased (**Fig. 5e** and **5f**) in the KO brain compared to the WT brain. These data indicate that KO of FOXO4 attenuates leukocyte infiltration.

**Figure 5.**
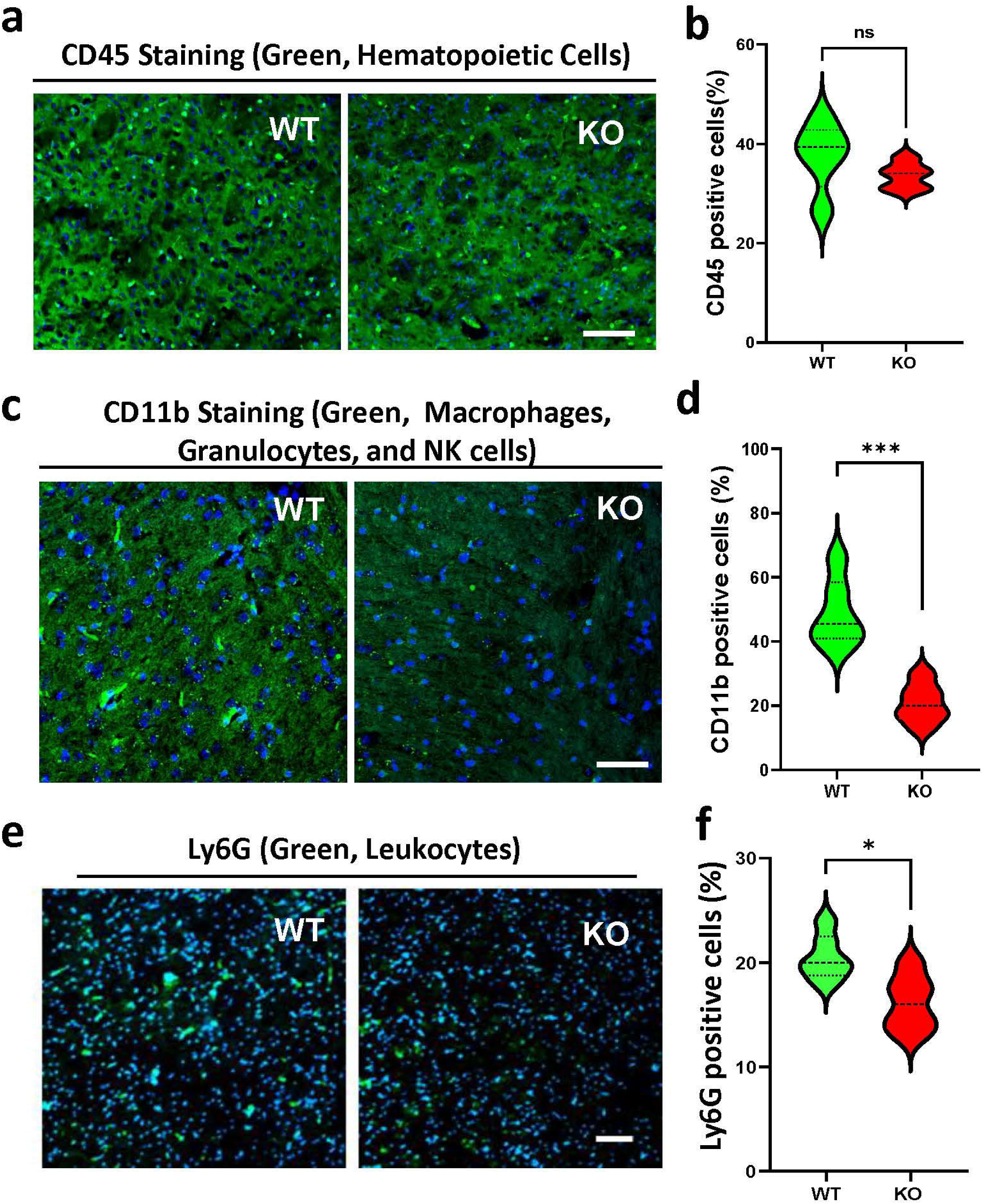
KO of FOXO4 attenuates leukocyte infiltration following I/R. **a.** Representative image showing CD45 staining. **b.** Quantitation of CD45-positively stained cells in the brain. **c.** Representative images showing CD11b staining. **d.** Quantitation of CD11b-positively stained cells. **e.** Representative image showing Ly6G staining. **f.** Quantitation of Ly6G-positively stained cells. Scale bar, 50 µm. Data are shown as mean ± SD; n = 5-7 for each group. ns, no significant difference; *p < 0.05, ***p < 0.001.

## Discussion

In this study, we investigated the role of FOXO4 in cerebral I/R-induced neuronal injury and validated that FOXO4 is a therapeutic target. While the protective role of FOXO4 in non-neuronal cells/tissues was previously reported *in vitro* and *in vivo* (27–29), to our knowledge, this is the first study of FOXO4 in I/R-induced neuronal injury in neurons and animal brains.

Unlike FOXO1 KO mice, which are embryonic lethal (30), FOXO4 KO mice do not differ from their WT littermates. Our findings revealed that KO of FOXO4 yielded a better outcome after cerebral I/R, significantly reducing the infarct volume 24 h after I/R, and improving functional recovery after that. KO of FOXO4 also enhanced learning and memory capability in mice after one week following the I/R. We also observed that the FOXO4 KO mice showed a significant decrease in inflammatory cytokines in the brain cortex 48 h after I/R. The inflammatory cytokines come from two sources: immune cells attracted from systemic circulation, and locally activated glial cells (31). Since we observed a significant decrease in CD11- and Ly6G-positive cells in the KO brain following I/R, leukocyte infiltration in the FOXO4 KO brain was reduced. Since CD11b is highly expressed by myeloid cells, the first line in the immune response (32), the significant decrease in CD11b in the FOXO4 KO brain should reflect reduced leukocyte infiltration through the microvessels.

Additionally, our results also support reduced activation of local microglia and astrocytes in the KO compared to the WT mouse brain. These findings are in good accordance with previous data showing the anti-inflammatory effect of deleting FOXO4 in an ischemic condition in another organ *in vivo* (33), and in cerebral endothelial cells in an *in vitro* model of I/R (27). After I/R, ROS, cell debris, mitochondrial dysfunction, excitotoxicity, and a disrupted blood-brain barrier (BBB) work together to trigger inflammatory responses, involving both local and systemic immune factors. As a transcription factor, FOXO4 regulates many pathological processes, such as apoptosis, endothelial function, ROS production, and BBB integrity, to facilitate cell death, whereas KO of FOXO4 leads to neuroprotective effects and impedes the progression of neuronal death and infarction, ultimately preserving functional performance, as supported by our findings.

In conclusion, using a genetically modified mouse, we validated FOXO4 as a therapeutic target for I/R-induced brain injury. Our data suggest that KO of FOXO4 reduces neuronal injury, promotes functional recovery, suppresses leukocyte infiltration, and attenuates neuroinflammation, indicating that FOXO4 is a potential therapeutic target to treat acute ischemic stroke caused neurological disorders.

## Acknowledgments

We would like to thank Drs. Yanying Liu and Christa Huber for their help in the early characterization of the mice. This work was supported by NIH/NINDS grant NS124846. Any opinions, findings, conclusions, or recommendations expressed in this material are those of the authors and do not necessarily reflect the views of the NIH.

